# Integrating Models to Decode the GnRH Pulse Generator

**DOI:** 10.64898/2026.01.05.697766

**Authors:** Kateryna Nechyporenko, Deyana Ivanova, Xiao Feng Li, Krasimira Tsaneva-Atanasova, Margaritis Voliotis, Kevin T. O’Byrne

## Abstract

The gonadotrophin-releasing hormone (GnRH) pulse generator is a critical neural oscillator that governs reproductive function through the pulsatile release of luteinising hormone (LH) and follicle stimulating hormone (FSH). Early electrophysiological studies, notably by Ernst Knobil, identified multiunit activity (MUA) volleys in the mediobasal hypothalamus that aligned with LH pulses, suggesting a neural basis for GnRH pulsatility. Although GnRH neurons exhibit some intrinsic secretory rhythmicity *in vitro*, their isolated electrophysiological signatures have proven inconsistent. Recent advances, including GCaMP fibre photometry in freely behaving mice, have revealed a precise correlation between episodic GnRH distal processes and LH pulses. However, it is now well established that a neural oscillator comprising hypothalamic kisspeptin neurones co-expressing neurokinin B and dynorphin, collectively termed the KNDy network, represents the core construct of the GnRH pulse generator. Understanding the dynamics of this network and its modulation by external inputs such as stress, metabolic cues, and circadian rhythms is essential. Computational modelling provides a systematic framework for integrating experimental data with mechanistic and predictive analyses to decode the GnRH pulse generator dynamics.

## Introduction

Successful reproduction depends on the proper function of a neural oscillator known as the gonadotrophin releasing hormone (GnRH) pulse generator, which drives the pulsatile secretion of the pituitary gonadotrophins, luteinising hormone (LH) and follicle stimulating hormone (FSH). These hormone pulses regulate folliculogenesis and ovulation in females and spermatogenesis in males. Initial attempts to elucidate the neural components of the GnRH pulse generator included the early work by Ernst Knobil and colleagues who discovered an electrophysiological correlate, namely multiunit activity (MUA) volleys recorded from the mediobasal hypothalamus of rhesus monkeys that faithfully aligned with LH pulses (1,2), in turn elicited by the pulsatile release of GnRH (3). The MUA volleys signature was subsequently shown in other species (4–6), but the identity of the neurones that compose them has remained enigmatic. It only seemed natural to focus on the GnRH neurones *per se*, and *in vitro* and *ex vivo* studies using immortalised cell lines and primary cultures of GnRH neurones, respectively, have indicated an ability of GnRH neurones to generate secretory pulses (7,8) implying an intrinsic pulsatile property of these neurones. However, electrophysiological recordings from single identified GnRH neurones *in vitro* have proved disappointing despite the occasional recording revealing a pattern of electrical activity that might reflect a neurosecretory burst underlying a pulse of GnRH secretion (9). Nevertheless, despite the recent demonstration of a perfect correlation between episodic increases in the activity of GnRH distal dendrons that terminate in the median eminence using, GCaMP fibre photometry, and LH pulses in freely behaving mice (10), we now know that the GnRH neurones are actually driven by an extrinsic neural oscillator comprising hypothalamic kisspeptin (Kiss1) neurones co-expressing neurokinin-B (NKB) and dynorphin (Dyn) (acronym-KNDy) (10–13). This KNDy network is now considered the elusive GnRH pulse generator neural construct and the focus of this review.

Through reciprocal excitatory and inhibitory interactions mediated by NKB and Dyn, respectively, KNDy neurones generate rhythmic bursts of activity that drive the episodic release of Kiss1 onto the distal processes of GnRH neurones approaching the median eminence (14,15), thereby synchronising pulsatile GnRH release. Even subtle disruptions to this finely tuned system can alter pulse frequency or amplitude, leading to reproductive disorders such as infertility. The GnRH pulse generator, however, does not function in isolation. Its activity is modulated by external inputs that convey information about physiological context and environmental conditions. Neural and hormonal signals from regions such as the paraventricular nucleus (16,17), medial amygdala (MeA) (18–22) and brainstem nuclei (23), along with peripheral cues related to stress (24), metabolic states (25,26), and circadian rhythms (27,28), modulate GnRH pulse generator excitability. These pathways enable reproductive function to adapt to changing conditions but also introduce additional layers of complexity that challenge experimental interpretation.

Mathematical modelling provides a powerful tool to unravel this complexity. Models of the GnRH pulse generator integrate experimental findings with theoretical frameworks to provide insight into mechanisms underlying rhythmic activity, support predictions of responses to perturbations, and link cellular interactions with physiological outcomes. In this review, we explore how the synergy between modelling and experimentation has transformed our understanding of the GnRH pulse generator. We trace the evolution of modelling alongside key experimental discoveries, highlighting how this integrative approach has helped to shed light on both the intrinsic oscillatory dynamics of the KNDy network and the modulatory influence of external inputs critical for control of reproduction.

### Temporal Signatures and Operational Characteristics of KNDy Neuronal Network

Over the years, modelling of the GnRH pulse generator has been carried out across multiple scales and levels of biological detail. Early phenomenological models linked fast GnRH neuronal activity with slower upstream regulation, later attributed to Kiss1 neurones (29–31). With increasing insight into the underlying physiology, more mechanistic single-cell models emerged. For instance, Lehnert and Khadra (32) incorporate autocrine GnRH feedback, calcium dynamics, and Kiss1-induced TRPC5 channel activation to investigate pulsatile neuronal activity and GnRH secretion. As the role of KNDy neurones became more established, population-level models were developed incorporating network-level mechanisms and predicting system responses to perturbations.

The first population level model of the KNDy network developed by Voliotis and colleagues in 2019 showed how reciprocal NKB excitation and Dyn inhibition generate rhythmic activity characteristic of relaxation oscillations (11). In this framework, fast NKB mediated positive feedback and glutamatergic excitation drive collective spiking, while slower Dyn accumulation terminates activity to produce self-sustained oscillations. Crucially, the study demonstrated that altering basal activity causes qualitative changes in network dynamics: sustained optogenetic stimulation between 1 and 15 Hz induced regular LH pulses in estrous mice, whereas frequencies outside this range failed to evoke pulsatility. Notably, 1 Hz photostimulation initiated pulsatile LH secretion and 5 Hz increased pulse frequency. Model simulations likewise showed that pulsatility emerges only within a defined excitability range and that modifying NKB or Dyn moves the system between silent, oscillatory and tonic regimes. Subsequent modelling by Voliotis and colleagues in 2021 (33) incorporated changes in network excitability across the ovarian cycle informed by experimental evidence that estradiol (E_2_) reduces NKB and dynorphin expression while enhancing glutamatergic signalling via increased vesicular glutamate transporter 2 expression. They showed that these steroid-dependent modulations shift the bifurcation boundaries defining the KNDy network’s pulsatile regime (33). These predictions align with experimental findings: low frequency optogenetic stimulation accelerates LH pulses in estrus, when network excitability is high, but slows or fails to initiate pulses in diestrus. Sex steroid regulation of pulse generator excitability is further supported by evidence that selective estrogen receptor-α (ERα) deletion in KNDy neurones enhances glutamatergic transmission and disrupts estrous cycles (34).

While these modelling studies established the network level principles of KNDy driven pulsatility, recent advances in single cell imaging have transformed the ability to examine the cellular mechanisms underlying KNDy activity. Building on fibre photometry studies in freely behaving mice that identified brief KNDy synchronisation events (SEs) as reliable precursors of LH pulses (35,36), miniendoscopic GRIN lens recordings revealed that individual KNDy neurones are recruited into tightly structured SEs that precede every pulse (12,37). Moore and colleagues (12) showed that each event exhibits a reproducible temporal pattern in which some neurones consistently activate earlier than others, giving rise to a proposed “leader-follower” organisation that challenges the classical view of KNDy neurones as a homogeneous population. In contrast, Han et al. (37) observed variable activation sequences across events in males, suggesting that fixed leader-follower ordering is not necessary for generating pulsatile dynamics. These divergent findings hinted at sex specific differences, which a recent comparative study supported: females exhibit faster and more regular SE intervals than males, whereas gonadectomy eliminates SE frequency differences but preserves subtle differences in SE interval distributions, indicating both steroid-dependent and steroid-independent sex divergence in pulse generator dynamics (38).

The dichotomy between these findings since then has been interrogated in the modelling study of Blanco et al. (39), testing whether temporal hierarchy as proposed by Moore et al. (12) is an intrinsic property of the KNDy neurones or emergent from network architecture. In the modelling framework they assume that the network is arranged into tightly connected neuronal clusters that are only weakly linked to one another. Within this modular architecture, their study shows that stable leader–follower ordering is present only when inter-cluster coupling is strong and intra-cluster coupling is relatively weak. Under these conditions, the most excitable cluster repeatedly initiates events, giving the appearance of dedicated leader cells, similar to the pattern observed *in vivo* by Moore et al. (12). In contrast, when intra-cluster coupling dominates, clusters become more autonomous, and as a result any cluster can initiate a synchronised event, making the firing order variable from one event to the other, which is consistent with the findings by Han et al. (37).

These results raise the question of how KNDy neurones coordinate their activity at the single-cell level. Recent work proposes a glutamate-two-transition model in which glutamatergic excitation initiates synchrony within the network, Dyn gates event occurrence, and NKB amplifies recruitment to full activation (37). At the cellular level, KNDy neurones exhibit high input resistance and burst prone dynamics shaped by I_h_, persistent Na^+^ and A type K^+^ (Kv4.2) currents, with variation in Kv4 kinetics influencing how individual neurones integrate into the network (40). A recent study showed that E_2_ reconfigures ionic conductances in ARC Kiss1 neurones, shifting activity from synchronous high-frequency firing to short burst firing (13). E_2_ increased the mRNA expression of voltage-gated Ca^2+^ channels, as well as HCN1/2 and KCNQ2, resulting in a rise in whole-cell Ca^2+^ current density without altering channel kinetics. At the same time, E_2_ decreased TRPC5 and GIRK2 expression, reducing the slow EPSP that sustains synchronous firing. Deletion of TRPC5 channels from KNDy neurones using CRISPR confirmed that these channels are required for generating the slow EPSP (13).

These findings motivated the construction of a detailed Hodgkin-Huxley-type model that reproduced the dynamical transitions under different E_2_ levels (13). The model showed that synchrony relies on TRPC5-mediated depolarisation balanced by GIRK inhibition, and that E_2_-driven increases in Ca^2+^ and I_h_ shift the system into burst firing optimised for glutamate release.

The above studies illustrate how the synergy between theoretical frameworks (Fig. 1(**A**)) and experiments (Fig. 1(**B**,**C**)) has enabled examination of KNDy neurone function across multiple scales, from population dynamics to network organisation and single cell mechanisms (Fig. 1(**A**)), thereby revealing the biological processes that shape their activity. However, our understanding remains incomplete, as the specific neuronal network “tipping-point” mechanisms that initiate and coordinate synchronised episodes (SEs) (41) of KNDy neuronal activity are not yet fully defined. Thus failing to recapitulate the synchronisation dynamics of the GnRH pulse generator. Apart from SEs that recruit the whole population, smaller “mini-synchronised events” (mSEs) occur in which only a subset of neurones elevate intracellular calcium (12,37). The mSEs are likely not to be accidental fluctuations but key intermediaries in the build-up toward a whole population SE (Fig. 1(**C**)), implying that mSEs may capture the earliest stages of network coordination. The observations by Han et al. (41) suggest that glutamate-dependent coupling between small groups of ARC Kiss1 neurones give rise to the mSEs. Complementing these findings, modular network model by Blanco et al. (39) demonstrates the partial firing of the network arises when a part of the said network reaches the excitability threshold, while the rest of the network remains below the basin of attraction associated with the state transition. The recently published single-cell-KNDy model (13) has been parametrised using an extensive set of electrophysiology data, that makes it a powerful tool in investigating the mechanisms that give rise to the mSEs. The model enables systematic exploration of how cell intrinsic properties, network topology and neuronal heterogeneity interact to produce partial recruitment events. In doing so, it provides a tool to identify the tipping point dynamics that orchestrate SEs, while teasing apart the contribution of the steroidal neuromodulation, for example from E_2_.

**Fig. 1:**
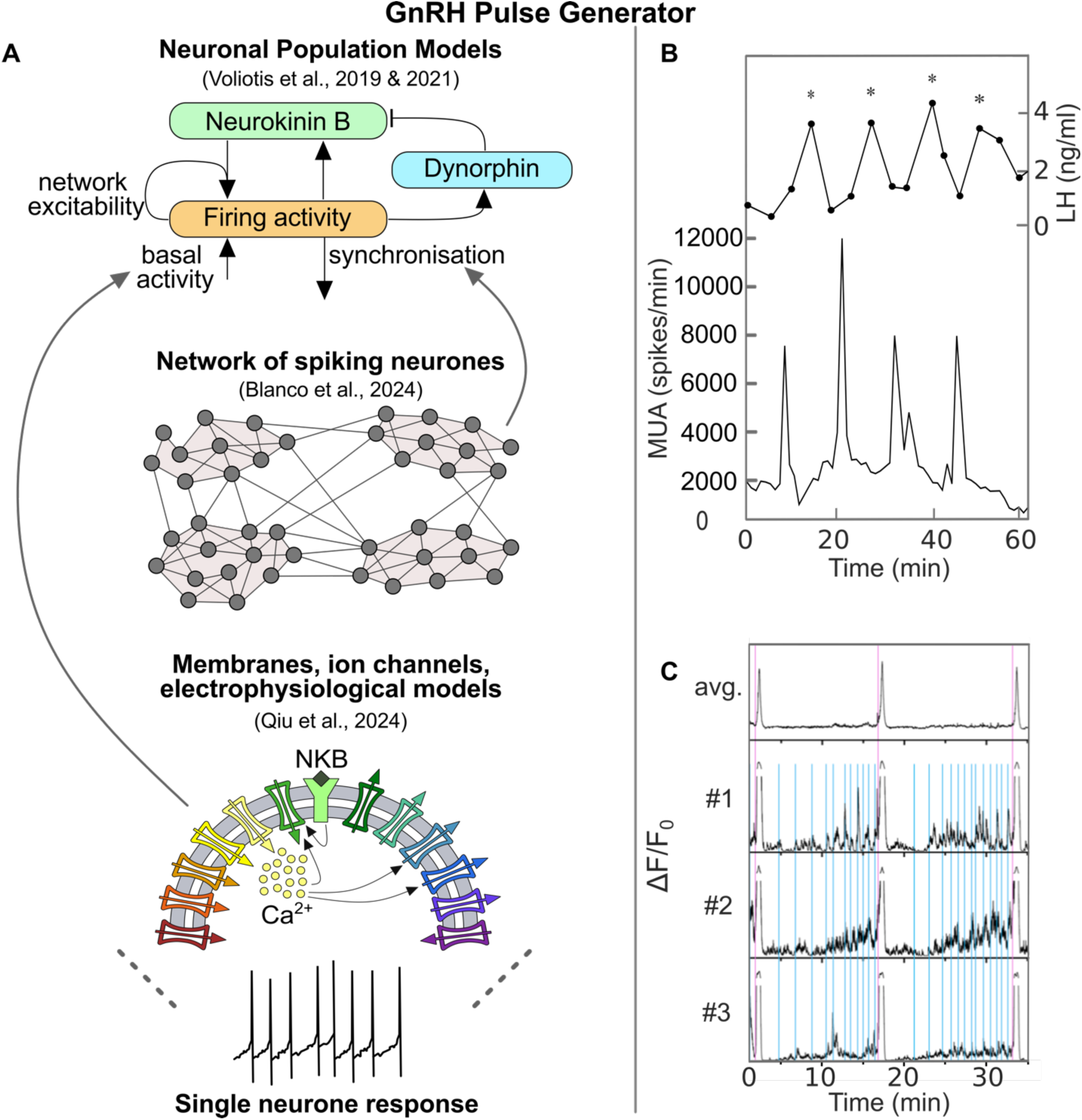
Integrating experimental and modelling frameworks to understand the GnRH pulse generator. (**A**) Schematic overview of mathematical modelling approaches across multiple scales, from neuronal population models (11,33) to spiking neuronal networks (39) to membrane and ion channel dynamics (13), shown from top to bottom. (**B**) MUA volley frequency recorded in the ARC of Kiss1Cre ovariectomised adult female mice, shown alongside corresponding LH pulses. (**C**) Representative *in vivo* GRIN lens miniendoscopic recording of calcium activity in KNDy neurones from diestrus mice. Top panel, population level KNDy dynamics showing synchronisation events (SEs) associated with pulsatile GnRH release. Three lower panels, representative single neurone calcium traces from three KNDy neurones within the field of view, illustrating a rich repertoire of mSEs (blue) in addition to full scale SEs (purple).

### External neuromodulation of the GnRH pulse generator

KNDy network intrinsic oscillatory behaviour is continually shaped by diverse excitatory and inhibitory inputs that fine tune their oscillatory rhythm to internal physiology and external influences. For instance, prenatal androgen exposure remodels upstream reproductive neural circuits in ways that elevate KNDy-driven LH pulsatility and impair steroid negative feedback, a mechanism now also implicated in human PCOS (42). Tracing studies show that KNDy neurones receive dense hypothalamic innervation from the ARC, paraventricular nucleus (PVN), dorsomedial hypothalamus (DMH) and the anteroventral periventricular nucleus (AVPV) populations conveying metabolic, circadian, stress and steroid feedback, alongside additional afferents from the limbic system, including septum, bed nucleus of the stria terminalis (BNST) and medial amygdala (18,43,44).

### Posterodorsal medial amygdala

Outside of the hypothalamus, the posterodorsal subnucleus of the medial amygdala (MePD) is a key limbic input to the GnRH pulse generator, relaying emotional, social, and stress-related information directly to KNDy neurones (19,20,22,45–47). Monosynaptic mapping indicates the presence of MePD to ARC projections (44), and a rapidly expanding body of experimental evidence and *in silico* modelling now shows that the MePD regulates the ARC KNDy network through coordinated Kiss1, glutamatergic, GABAergic, urocortin-3-corticotrophin-releasing factor type-2 receptor (UCN3-CRFR2) and neurokinin 3 receptor (NK3R) neuronal pathways.

Early work revealed that the MePD contains a small population of Kiss1 neurones and optogenetic stimulation of these neurones accelerates LH pulse frequency, establishing an excitatory drive from the medial amygdala to the GnRH pulse generator (19), expanding on earlier neuropharmacological studies (48). This contrasts with the amygdala’s broader inhibitory influence on reproduction (49), suggesting involvement of the other MePD circuit components in Kiss1’s pathway to KNDy. Glutamate and GABA are major excitatory and inhibitory neurotransmitters, respectively, and many neural networks rely on their balance to regulate their activity. Unsurprisingly, the presence of both has been identified in the MePD (50–52). To interrogate how GABA and glutamate signalling regulate MePD function, Lass et al. have used a combined approach of pharmacological antagonism of GABA or glutamate receptors with optogenetic stimulation of MePD Kiss1 neurones while measuring LH release (46). Although the inhibition of each of the neurotransmitters’ receptors alone produced no significant changes in the LH pulsatility, blocking MePD GABA receptors during Kiss1 photostimulation reversed the expected increase in LH pulse frequency. At the same time, the combination of glutamate receptor antagonism and stimulation of Kiss1 abolished the pulsatile release of LH.

To probe the underlying mechanisms, the established KNDy network model (11,33) was extended to incorporate model-derived MePD inputs, enabling investigation of how LH pulse frequency responds to antagonism of GABA and glutamate receptors in the MePD (46). The model of the MePD circuitry involves glutamatergic and GABA-GABA disinhibitory projections to the KNDy network, as well as glutamatergic activation of the GABAergic MePD projections, allowing to reproduce experimental results *in silico* (Fig. 2 (**A**)). The GABA-GABA disinhibitory pathway is consistent with the fact that MePD GABA is of pallidal origin (51). Additionally, Bian (53) defined three morphologically different subtypes of GABA neurones in the MePD. This may imply that these diverse populations contribute to distinct functional roles in MePD circuitry. The model (46) therefore suggests that MePD Kiss1 stimulation, when GABA receptors are blocked, lowers LH pulse frequency by driving glutamatergic activation of MePD GABAergic projections that target KNDy neurones. In other words, removing the ability of GABA interneurones to participate in disinhibition during optogenetic stimulation changes the MePD output from a net excitatory influence to a predominantly inhibitory one. Together, these modelling and experimental data identify MePD Kiss1 neurones as a modulatory “gain” on the GnRH pulse generator, acting through finely balanced glutamatergic and GABAergic neuronal circuits (Fig. 2(**A**)).

**Fig. 2:**
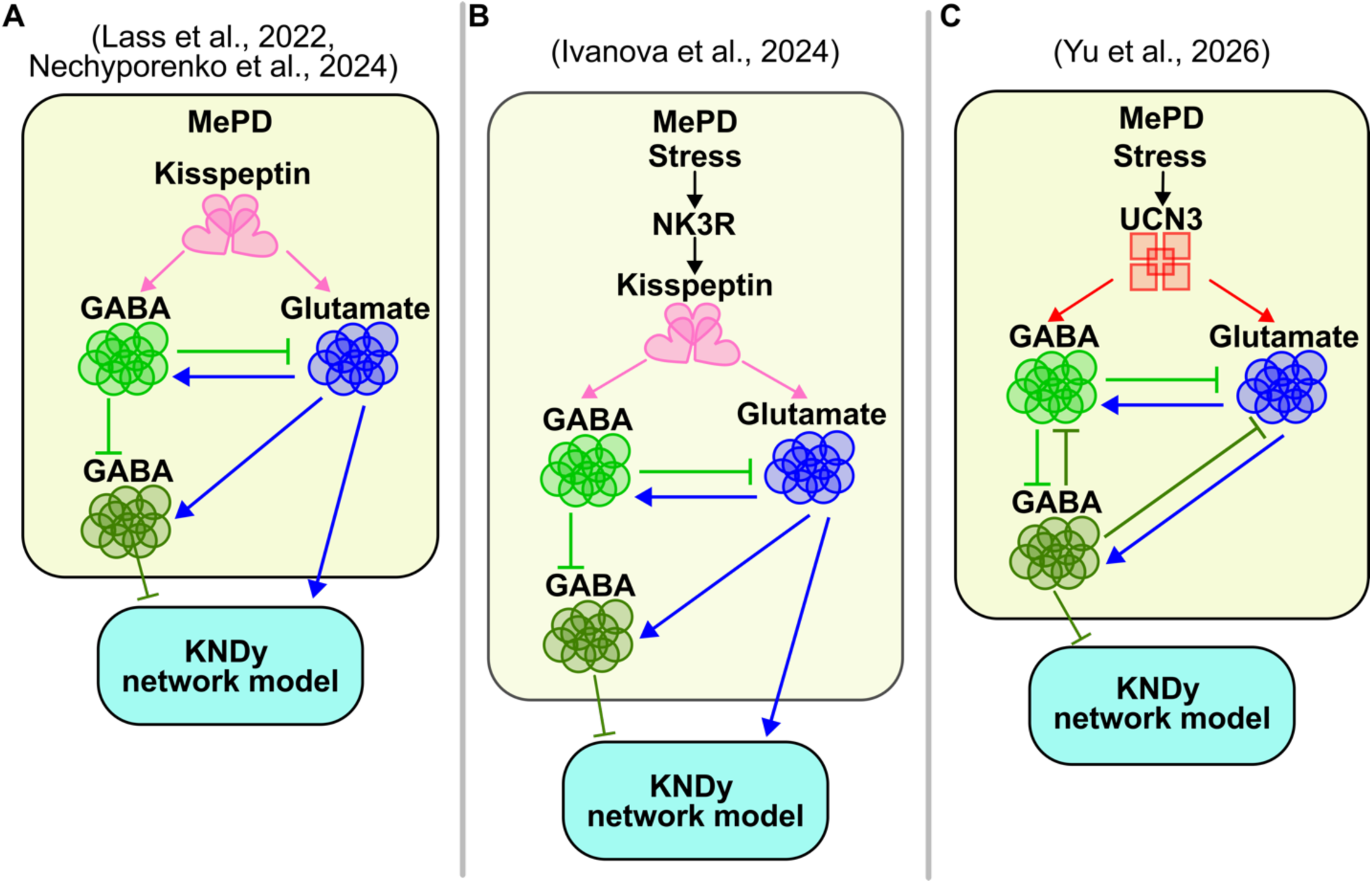
Schematic representation of the MePD circuit model used to study the effect of the interactions between the populations. (**A**) The modelling framework was first introduced by Lass et al. (46) where the ARC received a constant input. Nechyporenko et al. (22) subsequently examined the same framework to explore the emergence of oscillatory dynamics. (**B**) Ivanova et al. (54) extended this framework to incorporate the MePD NK3R pathway, demonstrating how kisspeptin-mediated signalling underlies predator odour induced stress inhibition of LH pulsatility. (**C**) Yu et al. (56) further used the MePD GABA-glutamate circuit to investigate the effects of UCN3 stimulation, treating this pathway as a mechanistic route through which stress modulates reproductive output.

The coupled MePD-KNDy model was then extended to integrate how MePD NK3R activity mediates predator-odour stress-induced inhibition of LH pulsatility in ovariectomised mice (54). The model predicted that stress (there predator odour) drives up the activity of MePD Kiss1 neurones by activating the NKB receptors expressed on these cells (Fig. 2(**B**)). Modelling in this study showed that NK3R activation increases MePD inhibitory output sufficiently to push the KNDy network above its upper oscillatory regime boundary, collapsing rhythmic activity. The MePD is shown to respond to physiological stressors (55), which makes it a good candidate to investigate the incorporation of stressors via the coupled model and study effects of stress on the GnRH pulse generator dynamics. In the model, stress is incorporated as an MePD NK3R-dependent input to the MePD GABA-glutamate circuit. Although NK3Rs are proposed to be expressed on MePD Kiss1 neurones and mediate stress effects through this neuronal population, they could also be present on other MePD neurones. The modelling framework therefore examines the functional consequences of NK3R-mediated modulation at the neuronal circuit level, without determining a specific cellular substrate through which stress ultimately shapes MePD output and, consequently, KNDy network dynamics.

The early coupled MePD-KNDy model treated MePD activity as a constant (averaged) input to the ARC, allowing the effects of GABA-GABA disinhibition and glutamatergic drive on KNDy network excitability to be examined (46,54). While these studies clarified how MePD Kiss1, GABA and tachykinin pathways drive the KNDy oscillator between active and silent regimes, in reality MePD circuit’s activity is likely more complex. Nechyporenko et al. (22) modelled MePD GABA and glutamate neuronal activity as oscillatory. The model analysis demonstrates that specific glutamate-GABA-GABA connectivity patterns determine MePD output polarity and, when coupled to the KNDy oscillator (Fig. 2(**A**)), reproduce the pharmacological effects previously reported by Lass et al. (46).

Importantly, the model predicted that stimulating MePD GABA projections suppresses LH pulses, consistent with *in vivo* optogenetic activation of MePD GABA terminals in the ARC (21). Additionally, the role of the glutamatergic projection was investigated, and the findings suggest that stimulation of the MePD glutamatergic projections has a non-monotonic effect on the GnRH pulse generator. This was confirmed experimentally, revealing a previously unrecognised excitatory MePD projection capable of modulating the KNDy network. Stimulation at 5 Hz increases LH pulse frequency, whereas higher frequencies (10 and 20 Hz) produce no significant change compared to basal (22). In the model, low levels of excitatory input enhance the activity of projection neurones, consistent with the 5 Hz experimental observation. With stronger excitation, mirroring higher stimulation frequencies, the system enters a regime where additional drive no longer yields proportional increases in activity. This behaviour suggests distinct modes of signal integration for GABA and glutamate in the MePD. GABAergic projections appear to convey unique, behaviourally relevant information from MePD to ARC, while glutamatergic projections contribute to homeostasis but do not encode specific instructive signals. This is consistent with the evidence showing that the majority of MePD projections are inhibitory (51–53). Therefore, increased glutamatergic drive is counterbalanced by local circuit mechanisms (22), an observation reflecting the largely inhibitory nature of MePD outputs.

The phenomenological nature of the MePD model (22,46,54) offers a flexible framework for testing the effects of other neuropeptides on circuit behaviour. Building on this, Yu and colleagues have used experimental and theoretical approaches to uncover how MePD GABAergic neurones relay stress signals to the reproductive axis (56). Using GRIN-lens mini-endoscopy, Yu et al. have recorded single-cell calcium activity, a proxy for neuronal activity, from MePD GABA neurones during both optogenetic stimulation of MePD UCN3 neurones and exposure to acute psychological stress (56). Clustering analysis has revealed two functionally distinct GABAergic subpopulations with anti-correlated activity patterns. Adapting the model of the MePD circuit (22) to describe input from UCN3 neuronal activity (Fig. 2(**C**)), it has been shown that anti-correlated activity is likely to arise from mutual inhibition between GABAergic populations. While it is possible that alternative connectivity patterns give rise to the anti-correlated activity in GABA populations, mutual inhibition naturally arises from the biophysical characteristics of GABAergic transmission, where the release of GABA inhibits target neurones, thereby producing inhibitory interactions between the subpopulations.

Reproducing anti-correlated activity between the two GABAergic subpopulations in the MePD, the coupled MePD-KNDy framework (56) successfully accounts for key experimental findings, including the effects of pharmacological blockade of MePD GABA and glutamate receptors with or without UCN3 stimulation (57), as well as the stress-induced loss of LH pulsatility (20). The model has been then used to predict how manipulating GABAergic signalling shapes the response to UCN3 input. It has indicated that activating MePD GABA neurones suppresses LH pulse frequency, whereas blocking GABA receptors prevents UCN3 stimulation from altering pulsatility. These predictions have been validated using an intersectional experimental strategy (56), supporting the modelling conclusions that MePD UCN3 influences the GnRH pulse generator primarily via a GABAergic pathway.

### Hypothalamic pituitary adrenal axis

There is growing evidence suggesting that the hypothalamic pituitary adrenal (HPA) axis, the core neuroendocrine system governing stress responses, has a dynamic interaction with the hypothalamic pituitary gonadal (HPG) axis, underpinning reproductive function (58,59). Stress-induced suppression of pulsatile LH secretion (17,59,60) highlights its role in modulating the GnRH pulse generator. Usually, the dynamic profiles of the HPG (11,33) and HPA (61) axes are considered as separate systems, which leads to a limited understanding of how the HPA and HPG axes coordinate their hormone rhythms and how perturbations to one will affect the other. To tackle this gap, a coupled-oscillator framework was introduced in which each oscillator represents a distinct aspect of neuroendocrine rhythmicity defined by its phase, frequency, and amplitude, enabling investigation of system-level outputs and cross-regulation between the stress and reproductive axes (28). The framework incorporates circadian rhythms, the glucocorticoid (CORT) oscillator, the GnRH pulse generator and the estrous cycle, making it possible to probe how stress input or E_2_ treatment at different cycle stages shapes pulse-generator dynamics. In particular, the model shows that high CORT levels increase the length of the estrous cycle and modulate response to acute stressors that have phase-dependent effects across the cycle. These findings can be interrogated through *in vivo* experimentation (28).

Within this context, the PVN is the primary hypothalamic driver of the HPA axis and a major locus for stress-induced modulation. Both optogenetic (17) and chemogenetic (62) activation of CRF neurones in the PVN have shown robust suppression of LH pulsatility. ARC Kiss1 neurones express CRF receptors and receive CRF neuronal projections (63), and activation of PVN CRF neurones exerts an inhibitory influence on the GnRH pulse generator resulting in decreased LH pulse frequency. Interestingly, PVN CRF neurones act indirectly through local GABA neurones within the PVN to suppress GnRH pulse generator frequency, as inhibiting PVN GABA neurones abolishes the inhibitory effect of CRF activation on LH pulsatility (17).

As was shown by Zavala and colleagues (28) the response to acute stress is cycle phase dependent. That prompts a question of whether activation of PVN CRF neurones has differential effects on LH pulsatility at different stages of the estrous cycle. Chemogenetic activation of PVN CRF neurones *in vivo* was shown to increase the length of the estrous cycle, specifically, the time spent in proestrus and diestrus was decreased, while the proportion of time spent in estrus and metestrus was significantly increased (64). Moreover, the LH frequency has decreased in metestrus, but unchanged in diestrus, confirming model predictions (64). The model (28) was adapted to include direct PVN output to the GnRH pulse generator and to CORT rhythms (Fig. 3), and re-calibrated to reproduce cycle periodicity in control animals. Under simulated CRF neuronal activation, the model captured the overall cycle lengthening and showed that although the absolute duration of diestrus is unchanged, it occupies a smaller fraction of the extended cycle (64). However, the predicted GnRH pulse generator frequency in metestrus was substantially lower than the experimental value, reflecting limitations of a phenomenological model where state variables represent oscillator properties rather than physiological quantities. The model also predicted an increase in CORT oscillation amplitude that would translate to a higher baseline. In contrast, the *in vivo* data have shown that morning CORT concentrations did not differ between the control and CRF-activated groups. If CORT was responsible for LH pulse suppression, elevated CORT would accompany reduced LH pulse frequency, yet this was not observed. This supports the interpretation that PVN CRF neurones suppress LH pulsatility independently of CORT, consistent with an indirect inhibitory pathway in which PVN CRF neurones signal through local PVN GABA neurones to inhibit ARC KNDy network activity (17).

**Fig. 3:**
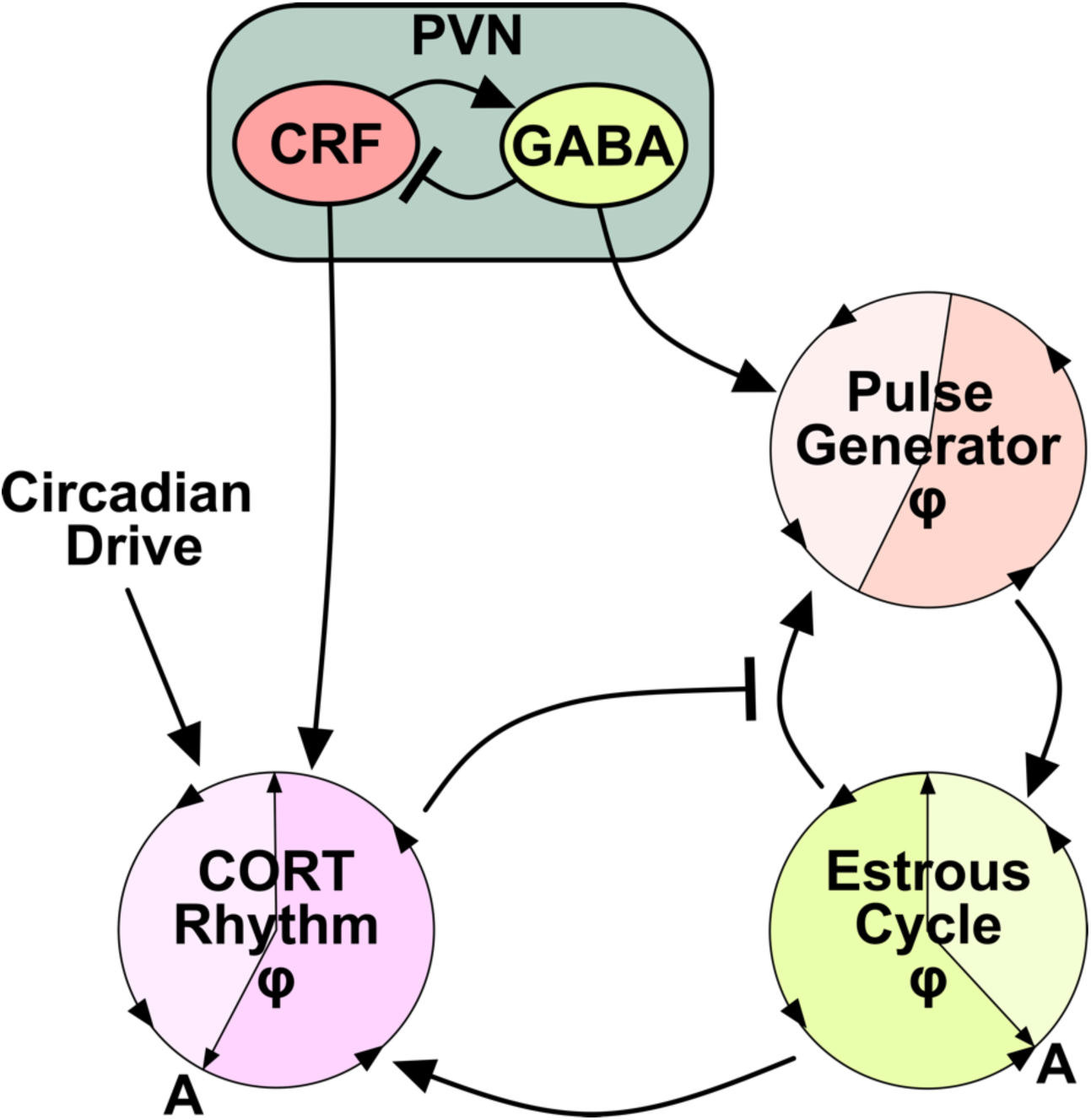
Network model of systems level cross-regulation between CORT rhythms, the hypothalamic GnRH pulse generator, and the estrous cycle, incorporating input from the PVN.

Together, these findings highlight that HPA-mediated suppression via adrenal output and PVN-driven suppression via local CRF-GABA circuits represent distinct mechanisms. To clarify how these pathways shape reproductive dynamics, it would be useful to develop a model of PVN circuitry that captures CRF-driven modulation of the GnRH pulse generator. Such a framework would allow systematic investigation of PVN contributions and provide deeper insight into how stress-related signalling shapes the temporal organisation of LH pulsatility.

## Future Directions

Recent advances in high-resolution calcium imaging provide a timely opportunity to further develop mechanistic models and improve our understanding of reproductive neuroendocrine control. At the level of the GnRH pulse generator, combining calcium imaging with detailed biophysical modelling offers a powerful route to understanding how SEs and mSEs arise. The recently developed single-cell KNDy model (13) provides a foundation for constructing network models composed of heterogeneous neurones with realistic ionic conductances. Coupling multiple virtual cells into a network model will enable systematic exploration of how intrinsic variability, population size and network architecture shape the emergence of mSEs, tipping-point dynamics and full population synchronisation, thereby linking single-cell properties to population-level activity.

Furthermore, systematic calibration of model dynamics to emerging MePD calcium recordings using likelihood-based or Bayesian inference will provide quantitative constraints on model parameters governing upstream modulatory processes. This integration will refine assumptions about underlying functional circuitry and synaptic interactions mediated by glutamatergic, GABAergic, and neuropeptidergic signalling; shedding light on the mechanisms essential for pulse generation and enabling a new round of experimentally testable model predictions.

Finally, future modeling efforts should incorporate the PVN-HPA axis into a unified, quantitative framework of stress-reproduction interactions. In such a model, the MePD would integrate fast, acute stress signals, the KNDy network would generate GnRH/LH pulses on intermediate timescales, and the PVN-HPA axis would provide slower hormonal feedback. This multiscale architecture would allow systematic testing of how stressors acting over different timescales gate reproductive output, and how acute neural inhibition and slower endocrine feedback interact to reshape the temporal organisation of GnRH and LH pulsatility.

## Acknowledgements

D.I., M.V., K.T.A., K.T.O., X.F.L. gratefully acknowledge the financial support of BBSRC via grants BB/W005913/1 (KCL), BB/W005883/1 (Exeter) and BB/S019979/1. KTA gratefully acknowledges the financial support of the EPSRC via grant EP/T017856/.

## Contributions

All authors contributed equally to writing the manuscript.

## Open Access

For the purpose of open access, the authors have applied a ‘Creative Commons Attribution (CC BY) licence to any Author Accepted Manuscript version arising from this submission.

